# Phylogenomic Analysis of Ants, Bees and Stinging Wasps: Improved Taxon Sampling Enhances Understanding of Hymenopteran Evolution

**DOI:** 10.1101/068957

**Authors:** Michael G. Branstetter, Bryan N. Danforth, James P. Pitts, Brant C. Faircloth, Philip S. Ward, Matthew L. Buffington, Michael W. Gates, Robert R. Kula, Seán G. Brady

## Abstract

The importance of taxon sampling in phylogenetic accuracy is a topic of active debate. We investigated the role of taxon sampling in causing incongruent results between two recent phylogenomic studies of stinging wasps (Hymenoptera: Aculeata), a diverse lineage that includes ants, bees and the majority of eusocial insects. Using target enrichment of ultraconserved element (UCE) loci, we assembled the largest aculeate phylogenomic data set to date, sampling 854 loci from 187 taxa, including 30 out of 31 aculeate families, and a diversity of parasitoid outgroups. We analyzed the complete matrix using multiple analytical approaches, and also performed a series of taxon inclusion/exclusion experiments, in which we analyzed taxon sets identical to and slightly modified from the previous phylogenomic studies. Our results provide a highly supported phylogeny for virtually all aculeate lineages sampled, supporting ants as sister to Apoidea (bees+apoid wasps), bees as sister to Philanthinae+Pemphredoninae (lineages within a paraphyletic Crabronidae), Melittidae as sister to remaining bees, and paraphyly of cuckoo wasps (Chrysidoidea). Our divergence dating analyses estimate ages for aculeate lineages in close concordance with the fossil record. Our analyses also demonstrate that outgroup choice and taxon evenness can fundamentally impact topology and clade support in phylogenomic inference.

## Introduction

The role of taxon sampling in improving phylogenetic accuracy is a topic of long-term controversy (1–9). Rosenberg & Kumar (8) argued that increasing the number of characters sampled is a better investment of resources compared to adding taxa. However, this conclusion has received much criticism and many subsequent studies have argued the opposite point (4,6), including some recent investigations that have employed genome-scale data (7,10,11). In the current age of phylogenomics, in which it is now possible to generate data sets with hundreds to thousands of loci (12,13), the argument over the relative importance of taxon versus character sampling has become largely irrelevant, with the more important question being: does improved taxon sampling increase phylogenetic accuracy? Here, we examine this question using a genome-scale data set that focuses on relationships within a major clade of insects.

Encompassing over 120,000 described species and having an estimated richness that might exceed two million species, the insect order Hymenoptera represents one of four insect megaradiations, (14–16). This extreme diversity includes many important lineages (*e.g.* sawflies, wood wasps, parasitic wasps), with arguably the most well known taxa belonging to the stinging wasps (Aculeata). The aculeates have attracted much attention because they include all eusocial Hymenoptera, most notably the ecologically and economically important ants and bees (17,18), and also the eusocial wasps (*e.g.,* paper wasps, hornets, and yellow jackets) (19). Eusociality has in fact evolved independently at least 6–8 times within the clade, making the group a model for studying the evolution of sociality (20–25). Outside of eusocial lineages, the group exhibits a wide range of life history strategies, with most species tending to be solitary or subsocial predators, specializing on a wide variety of arthropod prey (26,27). A number of taxa have also evolved endoparasitic or even herbivorous feeding strategies (*e.g.,* pollen and nectar) (14,15).

Given their diversity and importance, establishing a robust phylogeny and classification of the aculeates is of broad interest. Currently, the Aculeata includes over 70,000 described species and is divided into 9 superfamilies and 31 families (28). This classification is based upon a molecular study that found several morphologically circumscribed superfamilies and families to be non-monophyletic, most notably the Vespoidea, Bradynobaenidae, and Tiphiidae (28). More recent molecular studies have also provided new hypotheses for the phylogenetic positions of bees (29) and ants (30,31). Despite these improvements, considerable uncertainty exists as to the relationships among superfamilies and families within Aculeata.

To date, most molecular studies of Hymenoptera have used traditional Sanger sequencing methods, resulting in data sets with decent taxon sampling, but few loci and often low clade support (28,29,32–34). Several recent studies have instead employed next-generation sequencing approaches, but so far these have suffered from including few taxa (30,31,35). Two phylogenomic studies in particular produced conflicting relationships with regard to the phylogenetic position of ants. In the study of Johnson *et al.* (30) the authors used transcriptome data to resolve relationships among aculeate superfamilies and found ants to be sister to apoid wasps and bees (Apoidea), a novel and biologically attractive result. Conversely, Faircloth *et al.* (31), using hundreds of ultraconserved element loci (UCEs), recovered ants as sister to all other aculeate superfamilies (minus Chrysidoidea, which was not represented). Despite both studies employing genome-scale data, each produced highly supported but conflicting results. One potential problem for both studies was sparse taxon sampling, with Johnson *et al.* (30) including all superfamilies, but only 19 taxa, and Faircloth *et al.* (31) including 44 taxa, spanning six out of seven superfamilies, but with sampling biased towards the ants and missing a key outgroup (Chrysidoidea).

To test the hypothesis that taxon sampling caused the incongruent results between these phylogenomic studies, and to address important remaining uncertainties within Aculeata at the family level, we have generated the largest phylogenomic data set to date for the Aculeata. Building upon the study of Faircloth *et al.* (31), we have assembled a UCE data set comprising 187 taxa that includes all aculeate superfamilies, 30 out of 31 aculeate families (missing only Scolebythidae), and a diversity of outgroup superfamilies from across Hymenoptera. We analyzed the complete, 187-taxon matrix using multiple analytical approaches and recovered a highly supported phylogeny for virtually all aculeate lineages sampled. We also focused our sensitivity analyses on the placement of ants and bees within the Aculeata and found that taxon sampling can have a major impact on results even with genome-scale data.

## Results

### Sequencing Results

To generate our phylogenomic data set we used a recently developed approach that combines the targeted enrichment of ultraconserved element loci (UCEs) with multiplexed next-generation sequencing (36). We followed published lab protocols (31,36; see also materials and methods below) and used a Hymenoptera-specific probe set that targets 1,510 UCE loci from across the entire order. Using this approach we sequenced new molecular data for 139 taxa, and we combined these data with 16 taxa from Faircloth et al. (31) and 32 taxa from available genomes, resulting in a final data set that included 187 taxa (see electronic supporting information S1, Tables 1 and 2).

Within our taxon set we included 136 samples from within the Aculeata, representing 30 out of 31 recognized aculeate families (missing only Scolebythidae). Sampling within the Apoidea was particularly dense with 53 species sampled from 23 out of 25 recognized bee subfamilies, and 16 species from outside bees including the phylogenetically enigmatic families Ampulicidae and Heterogynaidae. We also included 14 species from four out of eight subfamilies within the paraphyletic family Crabronidae (29). For outgroup taxa, we sampled all superfamilies from within the sawfly grade (“Symphyta”), and 8 out of 12 non-aculeate superfamilies from within the Apocrita (“Parasitica”), including Trigonaloidea, Evanioidea, Ichneumonoidea, and Ceraphronoidea. Those taxa have been hypothesized in previous analyses as lineages closely related to Aculeata (15,32–34,37). To better compare results between the Johnson *et al.* (30) transcriptome study and our UCE study, we sampled DNA from 7 out of 12 of the same specimen series that were sampled in Johnson *et al.* (30).

After sequencing of enriched samples, we used the PHYLUCE v1.5 software package (38) to clean and assemble raw reads; extract, align and trim UCE loci (for sequenced and genome-enabled taxa); filter loci for taxon completeness, and generate DNA matrices ready for phylogenetic analysis (see materials and methods for details). For all taxa that we enriched and sequenced, we recovered an average of 966 UCE contigs per sample, with a mean contig length of 801 bp and an average coverage per UCE contig of 80X (for complete assembly stats see supporting information S1, Table 4). For genome-enabled taxa, we recovered an average of 1,036 UCE loci. Using our set of UCE alignments for all taxa, we evaluated the effects of filtering alignments for various levels of taxon occupancy (% of taxa required to be present in a given locus) and selected the 75% filtered locus set (“*Hym-187T-F75*”) as the primary locus set for analysis. The *Hym-187T-F75* locus set included 854 loci and had an average locus length of 238 bp resulting in a concatenated data matrix of 203,095 bp of which 143,608 sites were phylogenetically informative (for all matrix stats see supporting information S1, table 5).

### Phylogeny of Aculeata

After filtering for taxon completeness, we carried out maximum likelihood (ML) and Bayesian (BI) analyses on the concatenated *Hym-187T-F75* matrix using RAXML v8.0.3 and EXABAYES v1.4.1 (39), respectively. For both approaches we partitioned the data set using the kmeans algorithm available in a development version of PARTITIONFINDER (PF) (40), and for the ML searches we analyzed the matrix in several additional ways: (1) unpartitioned, (2) partitioned by locus, and (3) partitioned by the hcluster algorithm in PF v1.1.1 (data pre-partitioned by locus). We also ran three analyses using the summary method implemented in ASTRAL v4.8.0 for species tree estimation (41). For input into ASTRAL we generated bootstrapped gene trees for all loci using RAxML (200 reps). In the first analysis we used all individual gene trees and accompanying bootstrap trees as input into ASTRAL (854 loci total). In the second analysis we calculated and sorted loci by average bootstrap score (=informativeness) using R v3.2.2 (42) and we selected the 500 loci that had the highest scores for input into ASTRAL. We did this to reduce possible error/bias introduced by including uninformative loci, a problem that has been observed in other studies (43–45). For the third analysis we used all loci; however, to reduce error from loci with low information content we employed weighted statistical binning, which bins loci together based on shared statistical properties and then weights bins by the number of included loci (46) (details in electronic supporting material). We ran all species-tree analyses with 100 multi-locus bootstrap replicates (47).

To investigate other potential biases in our data, we carried out several additional analyses. In particular we wanted to address the observation that G+C variance can be a problem for reconstructing phylogeny in aculeate Hymenoptera (21). First, using PHYLUCE, we converted the complete, concatenated matrix to RY coding and we performed a best tree plus rapid bootstrapping analysis (100 bootstrap replicates) in RAxML using the BINGAMMA model of sequence evolution. Second, we filtered loci for various parameters calculated in R (scripts modified from (48)) and PHYLUCE: average bootstrap score, % invariant sites (= rate of evolution), and G+C variance. We then removed the 10% of loci that had the highest values for GC variance, and the top 10% of loci that had the lowest values for bootstrap score and % invariant sites. Following removal of outlier loci we retained 636 alignments (“best636”), and we concatenated these into a single matrix and analyzed the matrix unpartitioned in RAXML (best tree searches with 100 rapid bootstrap replicates). We did not partition the data because partitioning had little effect in the analysis of the complete matrix.

Across analyses we recovered a robust phylogeny of the Aculeata (Fig 1 and electronic supporting information S2, figures 1-14), with the topology being identical for all ML and BI analyses of the complete, non-RY-coded data, and nearly identical for the ST analyses and the ML analysis of the complete, RY-coded data (we recovered several differences within Chrysidoidea, noted below).

**Fig 1.**
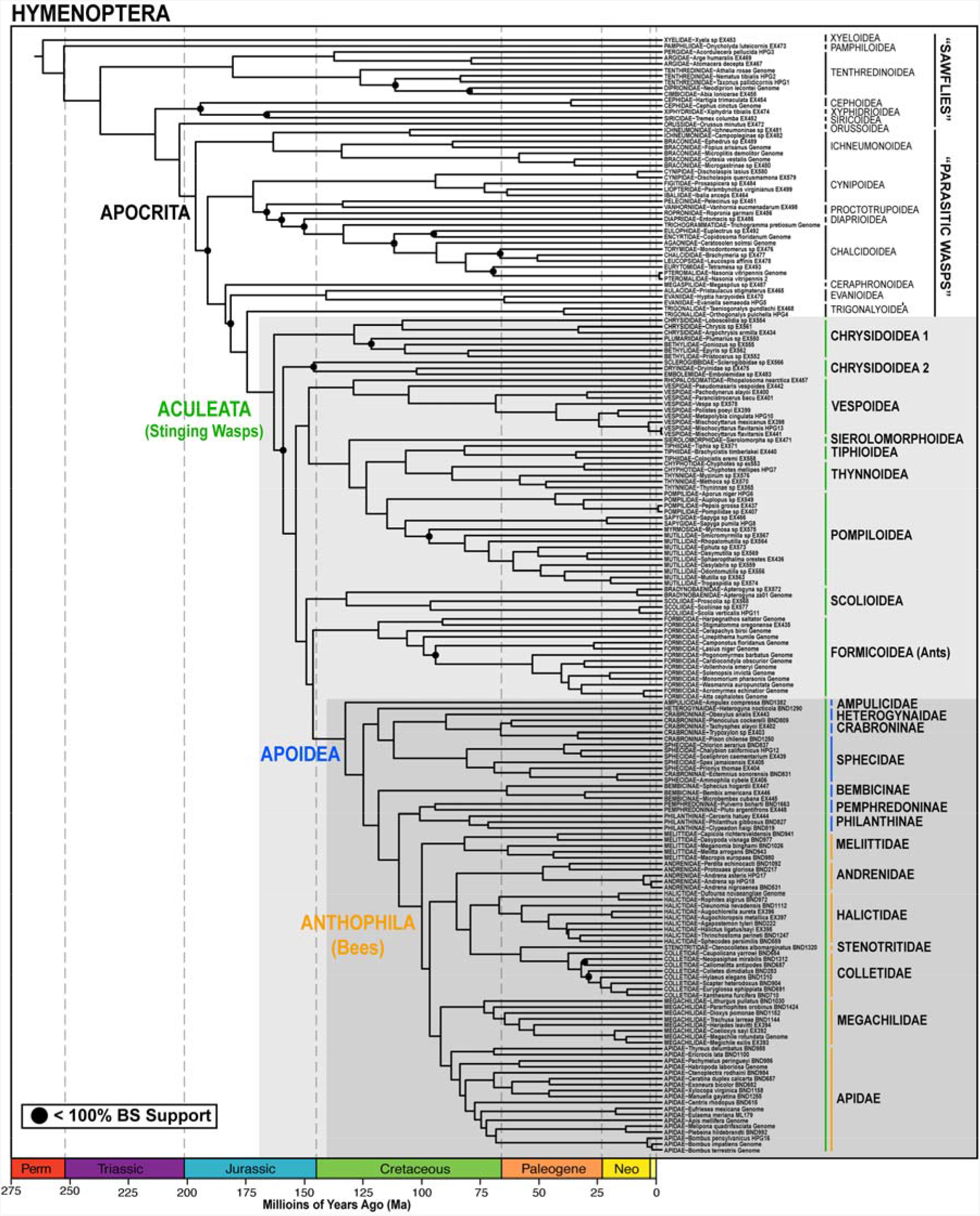
Dated phylogeny of Hymenoptera. We inferred the topology by analyzing the *Hym-187T-F75* matrix in RAxML (partitioned by kmeans algorithm; 854 loci; 203,095 bp of sequence data) and estimated the dates in BEAST (50 random loci; fixed topology; 38 calibration points). Black dots indicate nodes that received < 100% bootstrap support in the ML analysis.

We recovered the superfamily Trigonaloidea as sister to Aculeata in all analyses, and with maximum support except in the unbinned species tree analyses (97-98% support). Although we are missing several parasitoid superfamilies in our data set, this result is congruent with results from several recent molecular analyses (32,34,37), but is incongruent with results from (33). We did not recover the Ichneumonoidea, which has been a long-standing candidate as the sister group to the Aculeata (15), to be the sister group in any analysis. Within Aculeata, we recovered part of Chrysidoidea (cuckoo wasps and relatives) as sister to the remaining superfamilies, with Chrysidoidea itself paraphyletic, forming a grade of two (ML and BI analyses), three (binned ST analysis), or four (unbinned ST analyses) clades, depending on the analysis. In the ML and BI analyses of the non-RY-coded data, the first clade included [Chrysdidae+[Plumariidae+Bethylidae]] and the second clade included [Sclerogibbidae+[Embolemidae+Dryinidae]]. The placement of the second clade as sister to the remaining Aculeata, and the placement of Sclerogibbidae within the clade, received less than maximum bootstrap support in the ML analysis. In the analysis of the RY-coded data, we recovered a paraphyletic Chrysidoidea, but with only Sclerogibbidae falling outside of the superfamily. Results varied among the ST analyses, with the binned result being the same as in the non-RY-coded analyses except that Sclerogibbidae was placed outside of clade two and as sister to all remaining Aculeata. In the unbinned ST analyses the taxon *Plumarius* (Plumariidae) was moved out of the first clade mentioned above and placed as sister to Sclerogibbidae plus all other aculeates.

The remaining aculeate subfamilies separated into two major clades that were highly supported in all analyses. The first clade includes the superfamilies Vespoidea, Tiphioidea, Thynnoidea, and Pompiloidea, as well as the family Sierolomorphidae (currently in Tiphioidea). The monophyly of this group received maximum or nearly maximum support in all ML and BI analyses (≥ 98%), and slightly reduced support in the ST analyses (≥ 93%). Within the clade, we recovered a consistent topology across all analyses, with Vespoidea (includes Rhopalosomatidae and Vespidae) sister to the remaining superfamilies, and the phylogenetically enigmatic family Sierolomorphidae sister to [Pompiloidea+[Tiphioidea+Thynnoidea]]. Relationships among superfamilies received maximum support across analyses, except the monophyly of Vespoidea received less than maximum support in the ML analysis of the *best636* data set (98%) and the unbinned ST analyses (≥ 84%). Within Pompiloidea we recovered Pompilidae as sister to [Sapygidae+[Myrmosidae+Mutilidae]], but support for the position of Myrmosidae was less than maximum in all analyses except BI (≥ 57%), and support for the position of Sapygidae was reduced in a few analyses (≥ 74%), suggesting uncertainty. The second major clade contained the remaining aculeate superfamilies, with Scolioidea recovered as sister to Formicoidea+Apoidea in all analyses. This result received maximum support in all concatenated analyses. However, Scolioidea sister to Formicoidea+Apoidea received somewhat lower support in ST analyses (≥ 96%), and Formicoidea+Apoidea received 90% support in the binned ST analysis and only 43% and 7% support in the 500 best and all loci ST analyses. Overall, relationships among superfamilies largely agree with the recent Johnson *et al.* transcriptome study (30), except for the placement of Vespoidea.

Within Apoidea (bees and apoid wasps), our results are completely consistent across analyses and largely agree with Debevec *et al.* (29). We recovered Ampulicidae as sister to remaining taxa, and we found Crabronidae to be paraphyletic with respect to Sphecidae and bees. The remaining taxa formed a grade in the following order: [Heterogynaidae+[Crabroninae+Sphecidae], Bembicini, Phemphredoninae+Philanthinae, and the bees (Anthophila). The position of the enigmatic family Heterogynaidae as sister to Crabroninae+Sphecidae is a novel result, receiving less than maximum support only in the ST analyses (98% binned and ≥ 32% unbinned). The position of the bees as sister to the Pemphredoninae+Philanthinae was first reported in Debevec *et al.* (29) and was also recovered here with maximum support in all analyses except unbinned ST analyses (≥ 89%)

. Within bees, we recovered Melittidae to be sister to all remaining families, with maximum support in concatenated analyses (≥ 47% in ST analyses), as found in several previous studies (22). The remaining families were divided into two major clades: [Megachilidae+Apidae] (i.e., “long-tongued” bees *sensu* Michener (18)), and [Andrenidae+[[Stenotritidae+Colletidae]+Halictidae]]. Relationships of subfamilies within all families are largely congruent with previous studies of bee higher-level relationships (49). Within Apidae we recovered a monophyletic “cleptoparasitic clade” (50), monophyly of Anthophorini, Xylocopinae (51), and a sister-group relationship between Centridini and corbiculates. Relationships within corbiculates were notable because we recovered monophyly of the eusocial corbiculate tribes (Apini+Bombini+Meliponini) in all analyses except the ST analyses, which placed Apini as sister to [Euglossini+[Bombini+Meliponini]] with less than maximum support.

### Taxon Sampling Experiments

To test the effects of taxon sampling on phylogenetic inference and to examine the incongruent placement of ants between previous phylogenomic studies (30,31), we created and analyzed a series of alternative taxon sets, which can be divided into three categories (Fig 2): (1) variations of Johnson *et al.* (30), (2) variations of Faircloth *et al.* (31), and (3) variations of the current taxon set. For the first category, we generated two data sets, one with exactly the same taxon sampling as (30) (“*Johnson-19T*”), and one with the chrysidoid *Argochrysis armilla* removed (“*Johnson-18T*”). This particular manipulation was done because the major difference between the two phylogenomic studies was the presence/absence of Chrysidoidea, which is the sister taxon to the rest of Aculeata.

**Fig 2.**
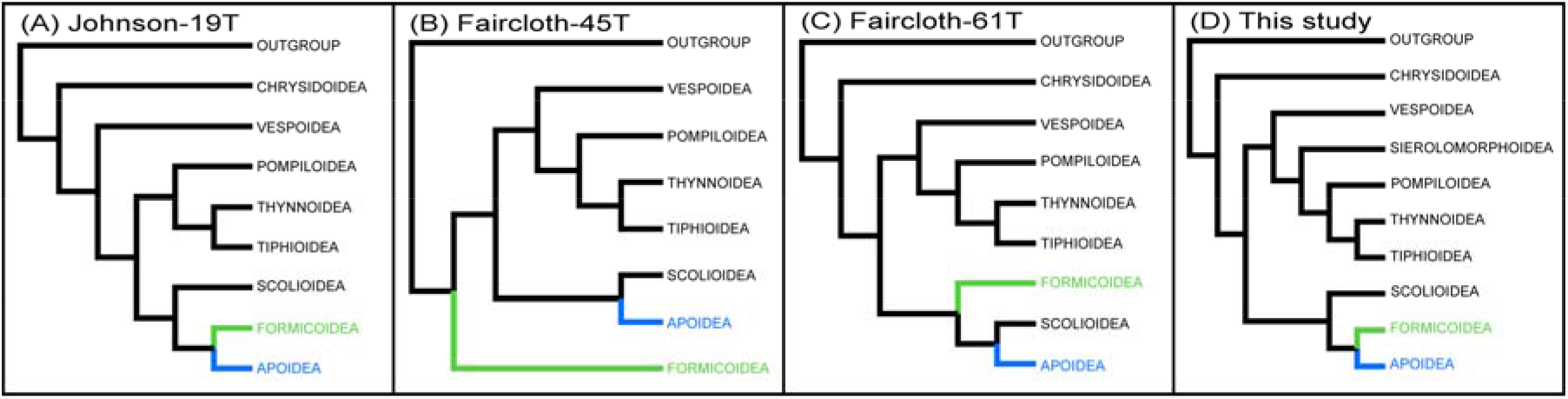
Alternative hypotheses for relationships among aculeate superfamilies. (A) Topology from Johnson *et al.* (30). (B) Topology from Faircloth *et al.* (31). (C) Topology from the *Faircltoh-61T* matrix analyzed in this study. (D) Preferred topology inferred in this study (includes Sierolomorphoidea). Topologies correspond to those reported in Table 1, except that topologies A and D are equivalent in terms of ants being sister toApoidea.

For the Faircloth *et al.* (31) manipulations we recreated the original 45 taxon matrix (“*Faircloth-45T*”) and then created several alternative taxon sets. First we added a single chrysidoid (“*Faircloth-46T*”), and then continued to add additional aculeates to balance the data set (“*Faircloth-52T*”, “*Faircloth-56T*” and “*Faircloth-61T*”). We also tried balancing the data set by removing most ant taxa from the original data set (“*Faircloth-26T*”) and adding in a chrysidoid (“*Faircloth-27T*”).

Finally, for the third category of taxon sampling experiments, we generated a data set with most outgroups removed (“*Hym-147T*”), leaving *Nasonia* as the earliest diverging outgroup and *Megaspilus* (Ceraphronoidea), Evanioidea, and Trigonaloidea as more recently diverging outgroups. From this taxon set, we removed chrysidoids (“*Hym-133T*”) and chrysidoids plus trigonaloids (“*Hym-131T*”). We also attempted to create the most balanced data set we could by removing excessive ant, bee and wasp taxa (“*Hym-100T*”). By removing distantly related outgroups, we not only reduced the number of taxa, but we potentially increased the average length of alignments. This is because UCE loci become more variable away from the central, core region (36) and alignment trimming (see materials and methods) removes poorly aligned regions. Thus, by removing more distant outgroups, alignments should be improve at the flanks of loci and less data should be trimmed.

In our description of the results we focus on the placement of ants (Formicoidea) among the other major lineages (superfamilies, etc.) of Aculeata. Among taxon sets, we recovered three alternative topologies (Fig 2, Table 1, and electronic supporting information S2, Figs 18-30): (A) ants sister to Apoidea, (B) ants sister to all other groups, minus Chrysidoidea, and (C) ants sister to Apoidea plus Scoliodea. In both of the Johnson *et al.* matrices, we recovered topology A. However, when we removed the chrysidoid, bootstrap support values for the relationships among ants, Apoidea, Scoliodea, Vespoidea, and Tiphiodea+Pompiloidea were reduced from maximum to 89%. We found a similar result in the analyses of the *Hym-147T* matrix and variants. All three matrices (*Hym-147T, Hym-133T,* and *Hym-131T*) produced topology A, but when chrysidoids and trigonaloids were removed (*Hym-131T*), support for the positions of ants as sister to Scolioidea+Apoidea was lowered to 90%.

**Table 1.**
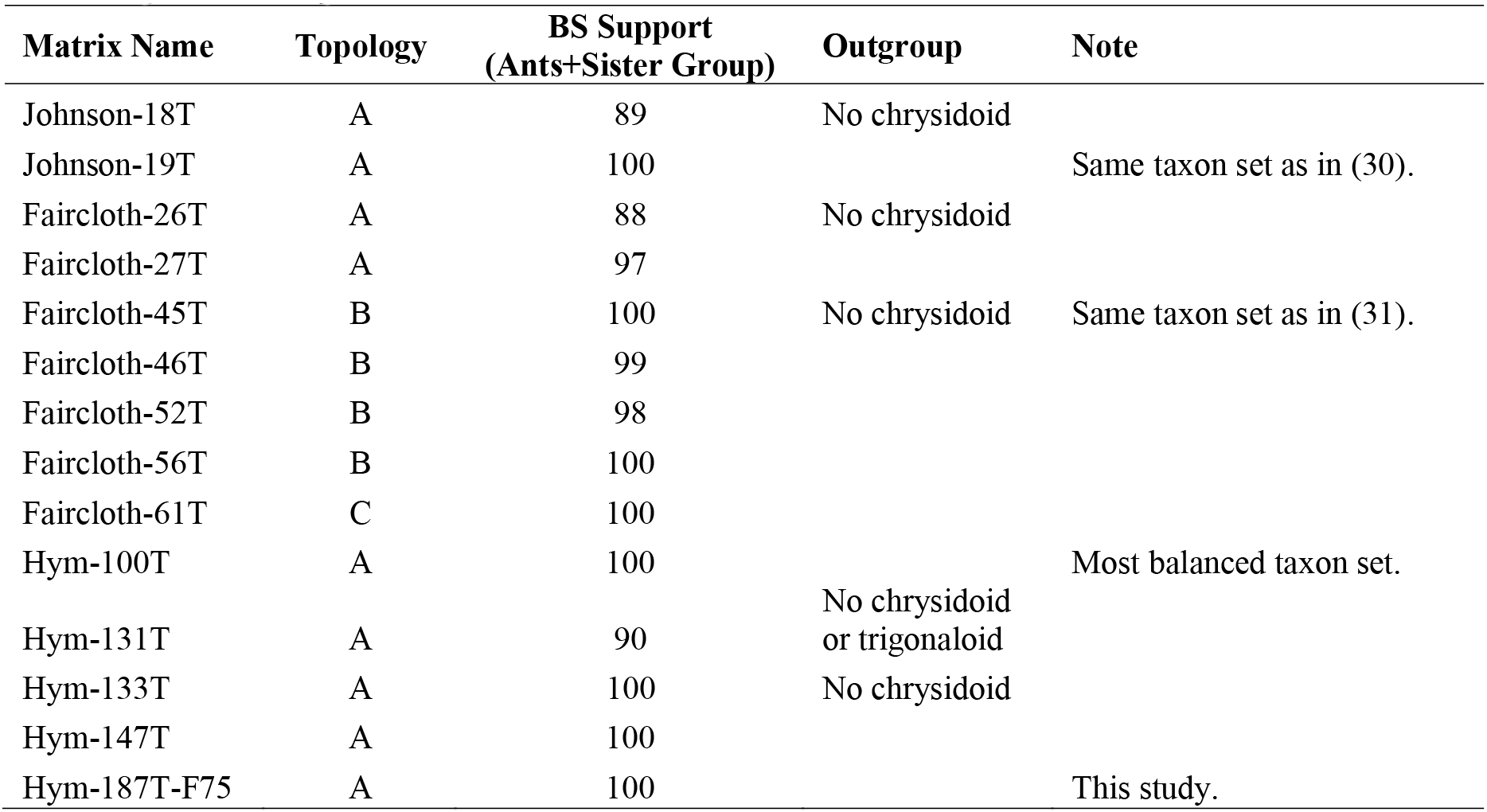
Results of the taxon inclusion/exclusion experiments as evidenced by topological and bootstrap support differences. The results suggest that both outgroup choice (chrysidoid presence/absence) and taxon evenness are important. The matrix name indicates whether the taxon set is a version of Johnson *et al.* (30), Faircloth *et al.* (31), or this study (“Hym”). Three different topologies were recovered: (A) ants sister to Apoidea; (B) ants sister to all other aculeate superfamilies, except Chrysidoidea; and (C) ants sister to Apoidea+Scoliodea. Bootstrap support indicates support for the clade that includes ants plus its sister group. Topologies correspond to those shown in Figure 1A-C, with regard to the position of ants.

Analysis of the original Faircloth *et al.* (31) taxon set (*Faircloth-45T*) produced topology B, as in the original study. Adding a chrysidoid to the taxon set (*Faircloth-46T*) did not change the topology, but did reduce support for the position of ants slightly (99%). In the *Faircloth-52T* and *Faircloth-56T* analyses, we also recovered topology B. However, in the *Faircloth-61T* analyses the topology shifted to C, placing ants as sister to Scolioidea plus Apoidea. The difference between *Faircloth-56T* and *Faircloth-61T* was the addition of several chrysidoids (Embolemidae and Dryinidae), Rhopalosomatidae (Vespoidea), and Ampulicidae (Apoidea), with the latter two taxa breaking long branches. Reducing and balancing the taxa of *Faircloth-45T* also altered the resulting topology. By reducing the number of ant taxa from 22 in *Faircloth-45T* to 3 taxa in *Faircloth-26T* the topology changed to A, but with only moderate support for Formicoidea+Apoidea (88%). Adding in a chrysidoid (*Faircloth-27T*) also resulted in topology A, and with nearly maximum bootstrap support for Formicoidea+Apoidea (97%).

Lastly, for the *Hym-100T* matrix, in which we reduced the number of ant and bee taxa to balance the larger taxon set, we recovered topology A, with the Formicoidea+Apoidea clade receiving maximum bootstrap support. All other relationships among superfamilies and within Apoidea were the same as those in the ML analysis of the *Hym-147T* and *Hym-187T* matrices.

### Divergence Dating

To generate a time tree for the evolution of the stinging wasps we estimated divergence dates for the complete 187 taxon matrix using the program BEAST v1.8.2 (52). We calibrated the analysis using 36 fossils representing taxa from across Hymenoptera and one secondary calibration taken from (53) for the root node (electronic supporting information S1, Table 3). For fossil ages we used midpoint dates taken from date ranges provided on the Fossilworks website (54) (http://fossilworks.org/). Due to computational challenges with BEAST, arising from having both a large number of taxa and a large amount of sequence data, we made the analysis feasible by inputting a starting tree (all nodes constrained), turning off tree-search operators, and using only a subset of the sequence data set rather than the entire concatenated matrix (details in electronic supporting material). We performed three separate analyses to compare the effects of different sets of loci on the final, dated results: (1) 25 loci that had the highest gene-tree bootstrap scores, (2) 50 loci that had the highest gene-tree bootstrap scores, and (3) 50 randomly selected loci.

The analysis of the three different locus sets (25 best loci, 50 best loci, 50 random loci) returned completely congruent dates (Table 2 and electronic supporting information S2, Figs 15-17). Consequently, we report here just the dates from the analysis of 50 random loci (Fig 2 and electronic supporting information S2, Fig 17). We estimated an age of 257 Ma (240-274 Ma 95% HPD) for crown Hymenoptera and 200 Ma (187-216 Ma) for Euhymenoptera (Orussoidea+Apocrita). The Apocrita arose 194 Ma (181-208), followed by the Aculeata at 161 Ma (154-169 Ma). Within Aculeata all of the superfamilies originated between 161 Ma to 100 Ma. The ants, minus the earliest diverging subfamilies Leptanillinae and Martialinae (not sampled in the current study), arose at least 118 Ma (108-128 Ma; Amblyoponinae+formicoid clade). The Apoidea arose 131 Ma (121-141 Ma), followed by the bees at 100 Ma (92-107 Ma).

**Table 2.**
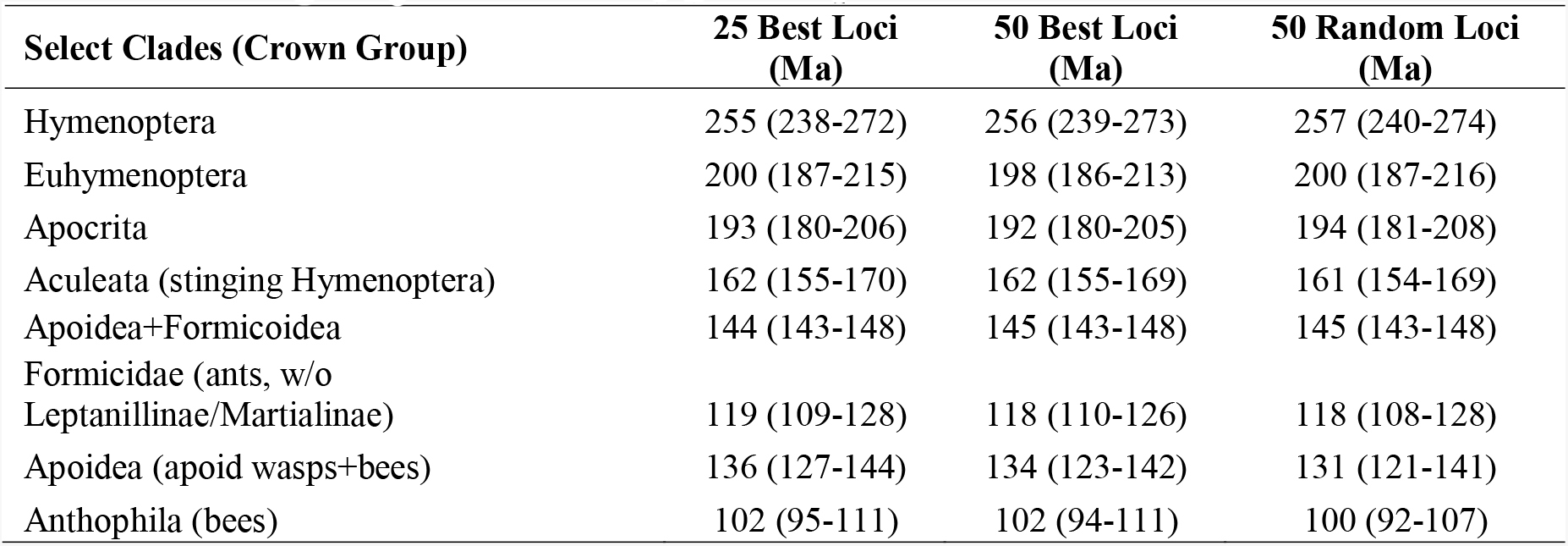
Divergence dates for key nodes (estimated with BEAST) comparing the 25 and 50 best loci (best equals loci with highest average gene-tree support values), and 50 randomly selected loci. Dates are given as median ages in millions of years ago (Ma), with the 95% highest posterior density given in parentheses.

## Discussion

The coupling of next-generation sequencing with reduced representation phylogenomics has driven a revolution in molecular systematics, making it possible to generate genome-scale data sets for hundreds of taxa at a fraction of the cost of traditional methods (12,13,55). Here, we further applied one of the most promising approaches, the target enrichment of ultraconserved elements (UCEs) (36), to the megadiverse insect order Hymenoptera, greatly extending a previous study which first introduced this approach in insects (31). We focused on family-level relationships of the stinging wasps (Aculeata) and produced a robust backbone phylogeny that confirms the utility of the UCE approach in Hymenoptera. In addition, by carrying out a series of taxon sampling experiments, we have demonstrated that even in the era of phylogenomics, careful taxon sampling and the use of taxon inclusion/exclusion experiments can be of critical importance.

Our phylogenomic results for Aculeata are largely consistent with and significantly amplify two previous molecular studies that employed traditional Sanger sequencing methods (28,29), and one recent transcriptome-based study (30). Compared to the two Sanger-based efforts, which both included a more limited number of taxa, our results agree in terms of the composition of superfamilies and families, with the only differences among studies being our finding that Chrysidoidea is paraphyletic and that the enigmatic family Sierolomorphidae is sister to [Pompiloidea+[Tiphioidea+Thynnoidea]]. The latter result supports resurrecting the superfamily Sierolomorphoidea, originally proposed by (56). Relationships among superfamilies, however, are quite different among these two studies, and our results mostly agree with those reported in the transcriptome study by Johnson *et al.* (30). An exception is the placement of Vespoidea in our study as sister to [Sierolomorphidae+[[Tiphioidea+Thynnoidea]+Pompiloidea]]]. This novel result is possibly due to our more extensive taxon sampling within Vespoidea (inclusion of Rhopalosomatidae) as compared to (30).

Our results strengthen previous findings of relationships within the Apoidea (29) and within the bees (Anthophila) (49). We confirm placement of Ampulicidae as sister to remaining Apoidea and the bees as sister to the crabronid subfamilies Philanthinae+Pemphredoninae. However, future studies should include an even broader sampling of Pemphredoninae and Philanthinae to confirm this hypothesis. Novel to our study is the placement of Heterogynaidae as sister to Crabroninae+Sphecidae. This taxon was previously placed as either sister to Apoidea (inside Ampulicidae) (29), sister to Astatinae+Bembicini (29), or sister to Philanthinae+Anthophila (57).

Within bees, our results provide further confirmation that the family Melittidae, previously thought to be sister to the long-tongued bees based on morphology (58), is monophyletic and sister to remaining bee families. It is also notable that most of our analyses recovered the eusocial corbiculate bees as monophyletic and sister to the weakly social Euglossini, thus favoring a single origin of eusociality within the group. Relationships among these taxa have been controversial, but our result agrees with a recent phylogenomic study that found that controlling for base-compositional heterogeneity, specifically GC variance among taxa, favored monophyly of eusocial corbiculates (21). The fact that we recovered this result without controlling for base compositional bias suggests that our UCE loci are robust to this problem, as was also suggested in another study of mammalian relationships (59).

Of major importance for understanding the evolution of eusociality, is our strongly supported result that Formicoidea (the ants) is sister to Apoidea (apoid wasps and bees). While disagreeing with the previous UCE study (31), it is in full agreement with the transcriptome study (30). The reason for the earlier conflict between these sources of phylogenomic data appears to be due to taxon sampling, with the earlier UCE study missing a key outgroup (Chrysidoidea) and having an excessive number of ant taxa (note that these were included intentionally to test the UCE method at resolving deep and shallow divergences), making the data set unbalanced. By conducting a series of taxon sampling experiments we demonstrated that excluding Chrysidoidea (or Chrysidoidea and Trigonaloidea) reduced bootstrap support for ants being sister to Apoidea. We also found that by either removing the disproportionate numbers of ant taxa, or adding additional taxa to the Faircloth *et al.* (31) taxon set, we were able to infer a topology consistent with both the transcriptome study and our more comprehensive taxon set presented here. Although the placement of ants as sister to Apoidea should still receive further investigation, we believe this result is the preferred one given its robustness across all of our analyses (ML, BI, and ST). Moreover, as discussed in Johnson *et al.* (30), the result is biologically attractive given that Apoidea includes the greatest number of eusocial Hymenoptera and all ants are eusocial. Furthermore, the finding that both bootstrap support and topology were affected by taxon sampling, provides additional evidence that taxon sampling in phylogenetics should still be a major concern, even in the age of phylogenomics, when data are no longer a limiting variable. Overcoming this challenge will require expanded and informed taxon selection as well as improved models and computational methods that can handle genome-scale data sets.

## Materials and Methods

### UCE Sequencing Pipeline

For all newly sampled taxa, we extracted DNA using Qiagen DNeasy Blood and Tissue kits (Qiagen Inc., Valencia, CA) and we fragmented up to 500 ng of input DNA to an average fragment distribution of 400-600 bp using a Qsonica Q800R sonicator (Qsonica LLC, Newton, CT). Following sonication, we constructed sequencing libraries using Kapa library preparation kits (Kapa Biosystems Inc., Wilmington, MA) and custom sample barcodes (60). We assessed success of library preparation following PCR amplification by measuring DNA concentration and visualizing libraries on an agarose gel. We purified reactions following PCR using 0.8 to 1.0X AMPure substitute (61).

For UCE enrichment we pooled 6–10 libraries together at equimolar concentrations and adjusted pool concentrations to 147 ng/µl. For each enrichment we used a total of 500 ng of DNA (3.4 μl each pool), and we performed enrichments using a custom RNA bait library developed for Hymenoptera (31) and synthesized by MYcroarray (MYcroarray, Ann Arbor, MI). The probe set includes 2,749 probes, targeting 1,510 UCE loci. We hybridized RNA bait libraries to sequencing libraries at 65°C for a period of 24 hours, and we enriched each pool following a standardized protocol (version 1.5; protocol available from http://ultraconserved.org).

We verified enrichment success with qPCR (ViiA 7, Applied Biosystems, Waltham MA) by comparing amplification profiles of unenriched to enriched pools using PCR primers designed from several UCE loci. After verification, we used qPCR to measure the DNA concentration of each pool, and we combined all pools together at equimolar ratios to produce a final pool-of-pools. To remove overly large and small fragments, we size-selected the final pools to a range of 300–800 bp using a Blue Pippin size selection instrument (Sage Science, Beverly, MA). We mailed size-selected pools to either the UCLA Neuroscience Genomics Core or the Cornell University Biotechnology Resource Center (http://www.biotech.cornell.edu/brc/genomics-facility), where the samples were quality checked on a Bioanalyzer (Agilent Technologies, Santa Clara, CA), quantified with qPCR, and sequenced on an Illumina HiSeq 2500 (2x150 Rapid Run; Ilumina Inc, San Diego, CA).

### Matrix Assembly

The sequencing facilities demultiplexed and converted raw data from BCl to FASTQ format using either BASESPACE or BCL2FASTQ (available at http://support.illumina.com/downloads/bcl2fastq_conversion_software_184.html). Using these files, we cleaned and trimmed raw reads using ILLUMIPROCESSOR (62), which is a wrapper program around TRIMMOMATIC (63,64). We performed all initial bioinformatics steps, including read cleaning, assembly, and alignment, using the software package PHYLUCE v1.5. For sequenced samples, we assembled reads *de novo* using a wrapper script around TRINITY v2013-02-25 (65). After assembly, we used PHYLUCE to identify individual UCE loci from the bulk of assembled contigs while removing potential paralogs. We then used PHYLUCE to combine the UCE contigs from the sequenced taxa with the contigs from the 32 genome-enabled taxa into a single FASTA file. We aligned all loci individually using a wrapper around MAFFT v7.130b (66), and we trimmed the alignments using a wrapper around GBLOCKS v0.91b (67,68), which we ran with reduced stringency settings (0.5, 0.5, 12, and 7 for b1–4 settings, respectively).

To extract an equivalent set of UCE loci from 32 genome-enabled taxa, we downloaded Hymenoptera genomes from NCBI and the Hymenoptera Genome Database (69). The genome of *Apterognya* za01 was provided by the authors of Johnson *et al.* (30). Using the software package PHYLUCE v1.5 (36,38), we aligned our UCE probe sequences to each genome and then sliced out matching sequence along with 400 bp of flanking DNA on either side (*i.e.*, 180 bp target plus 800 bp total flanking sequence). We then used the resulting UCE “contigs” for input into the downstream bioinformatics and matrix assembly steps.

### Analytical Details for Phylogenomic Inference

We investigated the tradeoff between taxon occupancy and locus occupancy (=missing data) in order to select a set of loci to be used for all remaining analyses. Using PHYLUCE, we filtered the entire set of trimmed alignments for different amounts of taxon completeness (% of taxa that must be included in a given alignment for it to be retained). This resulted in six locus sets filtered at a taxon threshold of 0, 25, 50, 75, 90, and 95% taxon completeness. To evaluate these locus sets we generated concatenated matrices and inferred maximum likelihood trees in RAXML v8.0.3 (70) (best tree search plus 100 rapid bootstrap replicates, GTR+Γ model of sequence evolution). We selected the best locus set by considering matrix completeness (more complete is better), topological consistency, and bootstrap support values (higher support is better). Using these criteria, we selected the 75% filtered set of alignments as the primary locus set for all subsequent analyses (electronic supporting information S1, Table 5; and S2, Figs 7-11).

All maximum likelihood (ML) analyses were performed using the best-tree plus rapid bootstrapping search (“-f a” option) in RAXML with 200 bootstrap reps for the kmeans analysis and 100 for all others. We used the GTR+Γ model of sequence evolution for all analyses (best tree and bootstrap searches). For the partitioned-Bayesian inference (BI) search, we executed two independent runs, each with four coupled chains (one cold and three heated chains). We linked branch lengths across partitions, and we ran each partitioned search for one million generations. We assessed burn-in, convergence among runs, and run performance by examining parameter files with the program TRACER v1.6.0 (71). We computed consensus trees using the *consense* utility, which comes as part of EXABAYES.

To carry out the weighted statistical binning ASTRAL analysis, we input all gene trees into the statistical binning pipeline using a support threshold of 75 (recommend for data sets with < 1000 loci). This grouped genes into 103 bins, comprising 73 bins of 8 loci and 30 bins of 9 loci. After binning we concatenated the genes into supergenes and used RAXML to infer supergene trees with bootstrap support (200 reps). We then input the resulting best trees, weighted by gene number, and the bootstrap trees, into ASTRAL and conducted a species tree analysis with 100 multi-locus bootstrap replicates (47).

For each taxon sampling experiment, we realigned the data after removing taxa, filtered alignments with GBLOCKS, filtered alignments for taxon completeness (using a 75% threshold), and generated a new concatenated matrix. We then analyzed each matrix in RAxML using a best tree plus rapid bootstrap search (100 replicates) with GTR+Γ as the model of sequence evolution.

As the input topology for the BEAST analyses, we used the best tree generated from the kmeans partitioned RAXML search of all loci. For each analysis, we concatenated the loci and analyzed the matrix without partitioning. We performed a total of four independent runs per analysis in BEAST, with each run progressing for 200 million generations, sampling every 1,000 generations. We also performed one search with the data removed so that the MCMC sampled from the prior distribution only. For the clock and substitution models, we selected uncorrelated lognormal and GTR+Γ, respectively. For the tree prior, we used a birth-death model, and for the ucld.mean prior, we used an exponential distribution with the mean set to 1.0 and the initial value set to 0.003 (determined empirically from preliminary runs).

## Acknowledgements

We would like to thank Dave Smith for donating specimens. We thank Jeffrey Sosa-Calvo, Ana Jesovnik, and Mike Lloyd for assistance with lab work. For sequencing we thank Joe DeYoung at the UCLA Neurosciences Genomics Core and Peter Schweitzer at the Cornell Genomics Facility. Lab work for this study was conducted at the Smithsonian NMNH Laboratory of Analytical Biology (LAB) and phylogenetic analyses were performed using the Smithsonian’s High-Performance Computer Cluster (Hydra) and the CIPRES Science Gateway. We thank X anonymous reviewers for helpful suggestions to the manuscript. Mention of trade names or commercial products in this publication is solely for the purpose of providing specific information and does not imply recommendation or endorsement by the USDA. USDA is an equal opportunity provider and employer.

